# Associations between Processing Speed and Psychopathology in a Transdiagnostic, Pediatric Sample

**DOI:** 10.1101/743021

**Authors:** Eliza Kramer, Bonhwang Koo, Anita Restrepo, Maki Koyama, Rebecca Neuhaus, Kenneth Pugh, Charissa Andreotti, Michael Milham

## Abstract

**Objective:** The present study sought to examine the relationships between processing speed (PS), mental health disorders, and learning disorders. Prior work has tended to explore relationships between PS deficits and individual diagnoses (i.e., anxiety, autism, ADHD, depressive) in isolation of one another, often relying on relatively modest sample sizes. In contrast, the present work simultaneously investigated associations between PS deficits and these diagnoses, along with specific learning disabilities (i.e., reading, math), in a large-scale, transdiagnostic, community self-referred sample.

**Method:** A total of 843 children, ages 8-16 were included from the Healthy Brain Network (HBN) Biobank. Given the presence of four PS tasks in HBN, principal component analysis (PCA) was employed to create a composite measure that represented the shared variance of the four PS tasks, referred to as PC1. Intraclass correlation coefficient (ICC) between the four PS measures, as well as PC1, were calculated to assess reliability. We then used multiple linear regression models to assess specific relationships between PS deficits and psychiatric diagnoses.

**Results:** ICCs were moderate between WISC-V tasks (0.663), and relatively modest between NIH Toolbox Pattern Comparison and other PS scales (0.14-0.27). Regression analyses revealed specific significant relationships between PS and reading and math disabilities, ADHD-inattentive type (ADHD-I), and ADHD-combined type (ADHD-C). Secondary analyses accounting for inattention dimensionally diminished associations with ADHD-C, but not ADHD-I or specific learning disability subtypes. The present study did not find a significant relationship with Autism Spectrum Disorder after accounting for inattentive symptoms. Consistent with prior work, demographic variables, including sex, socioeconomic status, and motor control exhibited independent relationships with PC1 as well.

**Discussion:** This study provided a comprehensive examination of PS, mental health disorders, and learning disabilities through a transdiagnostic approach. Implications for understanding how PS interacts with a highly heterogeneous childhood sample, as well as the need for increased focus on detection of affected populations are discussed.

## INTRODUCTION

Defined as the mental speed at which an individual is able to perceive and react to stimuli with reasonable accuracy^1, 2^, processing speed (PS) is a core component of human intelligence. In the context of early skill and knowledge acquisition, faster PS is thought to enable cognitive and academic progression by allowing for greater allocation of attention to higher-level tasks^3^. In other words, individuals with faster PS require less time and cognitive resources to carry out low level tasks, allowing for greater attention to more advanced cognitive developments. PS has been shown to be an important factor in academic achievement and test scores^4, 5^, as well as childhood peer relationships^6^ - indicating the importance of understanding the intricacies of PS deficits and their relations to function beyond academic performance. While impairments in PS are most commonly discussed in the context of learning and learning disorders, a growing literature has suggested potential links to a range of pediatric mental health disorders - thereby increasing the relevance of PS to the broader mental health community.

The relationship between PS and learning disorders is a largely studied topic, and the literature indicates that children with learning disorders frequently have PS deficits^7–14^. PS deficits have been most commonly associated with specific reading disorder (RD)^12, 15–17^, though one study found that the relationship between reading decoding and low PS disappears after the age of eight^3^. Though specific math disorder (MD) is less commonly studied with PS, some studies have found significant PS deficits in children with MD when compared to their typically-developing counterparts^11, 18, 19^, while others have found no significant differences in PS scores^20, 21^.

To date, beyond learning disorders, PS deficits are most consistently associated with attention-deficit/hyperactivity disorder (ADHD) in the mental health literature^27–38^. Some studies show that children with ADHD have lower PS scores than those with other clinical disorders, indicating PS has potential to help inform diagnosis^27, 28^. However, there is notable variation in severity of impairment associated with ADHD, mitigating this relationship^39^. In considering potential sources of variation across studies, it is worth noting mixed results for PS findings across ADHD presentations (i.e., inattentive (ADHD-I), hyperactive/impulsive (ADHD-H), combined (ADHD-C)). Some research has shown that all three presentations have similar PS deficits, indicating that PS is not a differentiating factor within ADHD presentations^29–32^. Others have found that ADHD-C and ADHD-I are both associated with PS deficits^28, 33^, while many have indicated that only children with ADHD-I demonstrated PS deficits^34–38, 40^. Of note, inattention has been specifically associated with PS deficits when treated as a dimensional symptom, as well as reading deficits^41–43^. Finally, it is worth noting that comorbidity amongst MD, RD, and ADHD has also been associated with low PS, and the relationships between these deficits continue to be explored^19, 22–26^.

PS deficits have been appreciated in mental health disorders beyond ADHD. Most notably, PS deficits have been explored in the pediatric autism spectrum disorder (ASD) literature^7, 8, 33, 44–46^, where specific associations with global functioning^47^ and general executive functioning deficits^7, 48–50^ have been recognized. A few studies have reported that PS deficits in ASD remain even after controlling for general intelligence^2, 51^, while other studies have failed to find any evidence of PS deficits either way^52, 53^. One study has attempted to explain ASD-related PS deficits in terms of motor demands^54^. The inconsistent results of PS findings in the ASD literature, similar to the observations in ADHD, make it difficult to understand the diagnostic usefulness of PS. Although less studied than ADHD and ASD, there is evidence of potential associations between PS deficits and internalizing disorders (e.g., anxiety, depressive). For example, a few studies found significantly lower PS scores for children with depression but not for those with anxiety^8, 55, 56^; though most of the literature has generally found non-significant relationships^7, 38, 57, 58^. Dimensional symptoms of depression and worry-related symptoms of anxiety were found to predict lower PS scores in a high school sample^59^.

The emerging literature suggesting associations between PS deficits and mental health disorders is not without limitations. Most notable is the tendency to look at specific disorders in isolation. It is well established that mental health disorders have a high degree of co-occurrence, especially between ADHD, ASD, SLD (specific learning disorder), and anxiety^15, 45, 53, 60–65^. Such comorbidities can confound the establishment of specific relationships when not appropriately accounted for; additionally, they may signal similarities in the underlying etiologies, which may be of interest. Compounding these challenges are variations in assessment strategies across studies, which can differ in the level of comprehensiveness of the diagnostic picture established for an individual, as well as whether they generate DSM categorical diagnoses or dimensional characterizations of psychopathology. These differences both hinder the ability to synthesize findings across studies, as well as decrease the likelihood of reproducing findings. Finally, consistent with the larger literature for pediatric mental health and learning is the reliance on relatively modest or moderate sample sizes, which inherently increases the risk of generating irreproducible findings.

In this study, using the large-scale, transdiagnostic sample being generated by the Child Mind Institute Healthy Brain Network (HBN) Biobank, we provide a comprehensive examination of associations between PS deficits and mental health disorders. Given that to date, there is no gold standard assessment of PS, we leverage the presence of the four assessments of PS in HBN (NIH Toolbox Pattern Comparison Task (NIH PC), Wechsler Intelligence Scale for Children, Fifth Edition (WISC-V) Symbol Search, WISC-V Coding, and a computerized adaptation of WISC-IV Symbol Search) to improve the robustness of our assessment of PS. Specifically, we used principal component analysis (PCA) to calculate a composite score from the four assessments for the purposes of our primary analyses. Additionally, we assess reliability among the four measures to provide insights into their comparability for future work. Finally, we employ a multiple regression framework to establish the specificity of associations between key mental health disorders (ADHD subtypes, SLD subtypes, ASD, anxiety, depression) and deficits in processing speed - a key piece of knowledge needed for efforts focused on establishing transdiagnostic perspectives of mental illness.

## METHODS

### Participants

Data were obtained from the Child Mind Institute Healthy Brain Network (HBN) Biobank (release 6.0; n = 2093), which uses a community self-referred recruitment model to generate a transdiagnostic sample; the sample is largely comprised of children (ages 5.0-21.0) affected by one or more mental health and learning disorders (Alexander et al., 2017). Children taking stimulant medication are asked to discontinue use while testing unless instructed by their physician to do otherwise. Research assistants obtained informed consent and assent before study tasks commence. There were two study sites for participants: one in Staten Island, NY and one in Midtown, NYC. We identified datasets to be included in the present work based upon the following criteria: 1) ages 8-16 (mean = 11.53 SD = 2.43); 2) complete data available for the four PS tasks (described below), 3) absence of extreme outlier scores (> 3 SD) for the PS tasks, 4) a full scale IQ score < 70.

### Processing Speed

A total of four tasks were used to estimate processing speed: Coding and Symbol Search which are two subtests of the WISC-V^66^ that make up the Processing Speed Index (PSI); a computerized adaptation of the WISC-IV^67^ Symbol Search task during the HBN EEG protocol (the WISC-IV was specifically chosen for this adaptation to avoid any repetition of stimuli with the WISC-V); and the NIH Toolbox^68^ Pattern Comparison task. See Table 1 for a brief description of each of these tasks.

**Table 1:** Processing Speed Task Descriptions

| **Task** | **Administrator:** | **Brief description** |
| --- | --- | --- |
| WISC-V Coding | Clinician | The Coding task requires participants to write a symbol that corresponds to a number 1-9, given a key at the top of the page. They have two minutes to complete as many as they can. Errors count negatively, omissions count neutrally |
| WISC-V Symbol Search | Clinician | The Symbol Search task requires participants to determine if one of two symbols has an identical match in an adjacent series of symbols. For every line of symbols, participants are required to mark the symbol that matches one of the two set aside on the left of the page. If the symbol is not repeated in the line, they mark the word "NO". They have two minutes to complete as many as they can. |
| WISC-IV Symbol Search Computerized | Computerized text instructions, RA to assist | Like the interviewer-based WISC-IV symbol search, each item contains two symbols and an adjacent series of symbols. However, children click "YES" if one of the two symbols is repeated, or "NO" if neither symbol re-appears in the series. They are instructed to complete as many items as possible in two minutes. |
| NIH Toolbox- Pattern Comparison | Computerized verbal instructions, RA to assist | Participants determine whether two images on the iPad screen are the same. Images differ by color, number of items, or completeness. Participants click YES if they are the same, and NO if they are not, as quickly and accurately as they can. Instructions are given verbally from the iPad with a trained research assistant in the room. |

### General Intelligence

All HBN participants were administered the core 10 subtests from the WISC-V by a licensed clinician to obtain a full-scale IQ (FSIQ), along with its component scores. WISC-GAI (general ability index) was also calculated. WISC-GAI is a measure of intelligence derived from weighted core verbal comprehension, fluid reasoning, and visual-spatial subtests, creating a measure of intelligence with a reduced emphasis on working memory and processing speed.

### Psychiatric diagnosis

During the last visit of the HBN protocol, the clinician administers the Kiddie Schedule for Affective Disorders and Schizophrenia (K-SADS), a semi-structured DSM-V-based psychiatric interview used to derive clinical diagnosis, to both the participant and the parent. The clinical team, including the licensed clinician, social worker (or junior psychologist), and psychiatrist (if consultation is needed), uses the K-SADS along with observations from clinical visits and questionnaires to provide a consensus diagnosis for each participant. The anxiety group in analyses includes children who received a current diagnosis of any anxiety disorder. The depression group in analyses includes children who received a current diagnosis classified as a depressive disorder, including major depressive disorder, persistent depressive disorder (dysthymia), disruptive mood dysregulation disorder, and other specified depression.

### Learning Disabilities

Specific learning disorder diagnoses were made by licensed psychologists based upon the reported educational history as well as results of the WISC-V, Wechsler Individual Achievement Test, Third Edition (WIAT-III), Test of Word Reading Efficiency, Second Edition (TOWRE-II), and relevant subsections of the Comprehensive Test of Phonological Processing, Second Edition (CTOPP-2). Given that these diagnoses were based on clinical judgment rather than a specific research cutoff, one concern is that they may be overly conservative relative to more commonly used research definitions for learning disability in the literature. To address this concern, we expanded our learning disabilities groups to additionally include any participant with a WIAT-Word Reading score < 85 for Specific Learning Disability-Reading (called the SLD-Reading group), and a WIAT Numerical Operations score < 85 for Specific Learning Disability-Math (called the SLD-Math group). Categorical regression analyses included these groups based on the above criteria.

### Parent and self-report questionnaires

Families were administered questionnaires to report on participants’ behavioral, social, cognitive, and emotional functioning. Parents completed questionnaires pertaining to their children on the computer while in the office. Participants electronically completed self-report questionnaires about themselves and their family while in the office. If a participant could not comprehend the questions or had difficulty with reading comprehension, a research assistant read the questions to the participant. The HBN includes the following relevant questionnaires: Demographics, Barratt (Scale for Socioeconomic Status)^69^, Child Behavior Checklist (CBCL)^70^, Screen for Child Anxiety Related Disorders (SCARED) parent and self-report^71^, Mood & Feelings Questionnaire (MFQ), parent and self-report^72^, Affective Reactivity Index (ARI) parent and self-report^73^, Strengths and Weaknesses Assessment of Normal Behavior (SWAN) parent report^74^, Autism Spectrum Screening Questionnaire (ASSQ)^75^, Social Communication Questionnaire (SCQ)^76^, and Social Responsiveness Scale-2 (SRS-2)^77^.

### Motor control

To control for possible confounds related to motor skills, the Grooved Pegboard Test (GPT)^78^ is used to assess eye–hand coordination and motor speed. It is critical to include motor control in studies related to processing speed due to their common overlapping deficit^34, 35, 46, 54, 60, 79^.

### Quality assurance

Intelligence testing is scored by the licensed clinician. Assessments are double scored by trained research assistants to ensure correct counting. Scores are then entered by a research assistant and double entered by a separate research assistant to ensure correct entry.

### Data Reduction

Principal components based on the four PS measures (WISC-V Coding, WISC-V Symbol Search (WISC-SS Clinician), WISC-IV Symbol Search (WISC-SS EEG), and NIH Pattern Comparison (NIH-PC)) were generated using the *prcomp* function in R. The first component from this analysis is used in these analyses and referred to as PC1, and the second component is referred to as PC2. The four measures’ raw scores were converted into standardized scores in order to run the PCA analysis, as the raw scores are on differential scales. Primary analyses in the present work were focused on PC1, which represented the common variance among the four tools (supplementary materials report secondary findings related to PC2).

### Statistical Analysis

Pearson’s correlation coefficient and single-fixed raters intraclass correlation coefficient (ICC) between the four PS measures and PC1 were calculated. In addition, Pearson’s correlation coefficient was calculated between PC1 and WISC-V FSIQ, WISC-V GAI, and the grooved pegboard dominant hand z-score to assess the relationship between PS, intelligence and motor control.

In order to assess the effects of psychopathology (i.e., categorical diagnoses) on PS, multiple linear regression analyses were performed on PC1, using categorical diagnoses from the clinician consensus diagnosis (ADHD-I, ADHD-C, ASD, anxiety, depression), the learning disability groups described above, as well as sex, age, SES, collection site, and grooved pegboard z-score as predictors. We also show results of each of these individual diagnoses in a regression with confounding variables predicting PC1, in order to determine how they relate to PC1 before accounting for the other diagnoses. The same analyses were run on each of the four PS subtests for comparison, yielding largely congruent results which can be found in the supplemental materials. The ADHD-Hyperactive presentation diagnosis was not included in the analysis due to the low number of participants with the disorder. A separate categorical regression that included WISC-V FSIQ was included to assess the role of overall intelligence in these relationships.

In addition, we examined the role of psychiatric symptomatology (i.e. symptom dimensions) in PS using Pearson correlations. Age and sex were regressed out of non-standardized symptomatology questionnaire scores. The scales that had a significant relationship with PC1 were then entered into a multiple linear regression model in order to examine how these dimensional symptoms interact with PS when accounting for each other. We entered the following scales into the regression to analyze their predictive value for PS: SWAN (Inattentive and Hyperactivity subscores), SRS, SCQ, ASSQ, sex, age, SES, collection site, and grooved pegboard dominant hand z-score. As with the categorical regression, dimensional symptoms were also run individually with confounding variables predicting PC1 to examine individual relationships.

Given observed patterns of relationship between ADHD presentations and PS, a post-hoc analysis of the impact of inattentive symptoms on the relationship with PS and clinician diagnosis was calculated in a multiple linear regression with SWAN-Inattentive scale as an added predictor.

## RESULTS

### Sample Characteristics

After participant exclusion, the total sample consisted of 843 participants (535 male, 308 female). 48.75% identified as white, 12.57% as black, 10.44% as Hispanic/Latino, 17.08% as two or more races, 6.29% as unknown/unavailable, 3.20% as Asian/Pacific Islander, and 1.67% as other. For 47.81% of children, one or both parents had a professional or managerial occupation. Regarding those excluded (n = 1,250), 828 were excluded due to age, 414 were excluded due to incomplete data (e.g., failure to complete the HBN EEG protocol, which included missing either the computerized adaption of WISC IV Symbol Search [n = 263]; one of the other PS tasks [n = 117]; incomplete demographics questionnaire [n = 34]), 5 were excluded due to having a score > 3 SD from the mean on one of the four PS tasks, and 3 were excluded due to completing the tasks at an off-site location.

Refer to Figure 1 for sample characteristics (age, sex, SES) for the most frequent categorical diagnoses in HBN, as well as information regarding diagnostic comorbidities. The two SLD subgroups were based on the larger definition of SLD described above. The total sample size per disorder is indicated by the horizontal bar that extends to the left of the diagnoses. The bars that extend vertically are the sample sizes that coincide with the diagnoses that have the black dot below. If there is more than one black dot with a line, that indicates comorbidity between the two or more disorders. The blue vs. red vertical bars indicate males and females, respectively. Below each dot(s) indicates the age and SES intervals for said diagnostic group. This figure was created using the package UpSetR within RStudio^80^.

**Figure 1.**
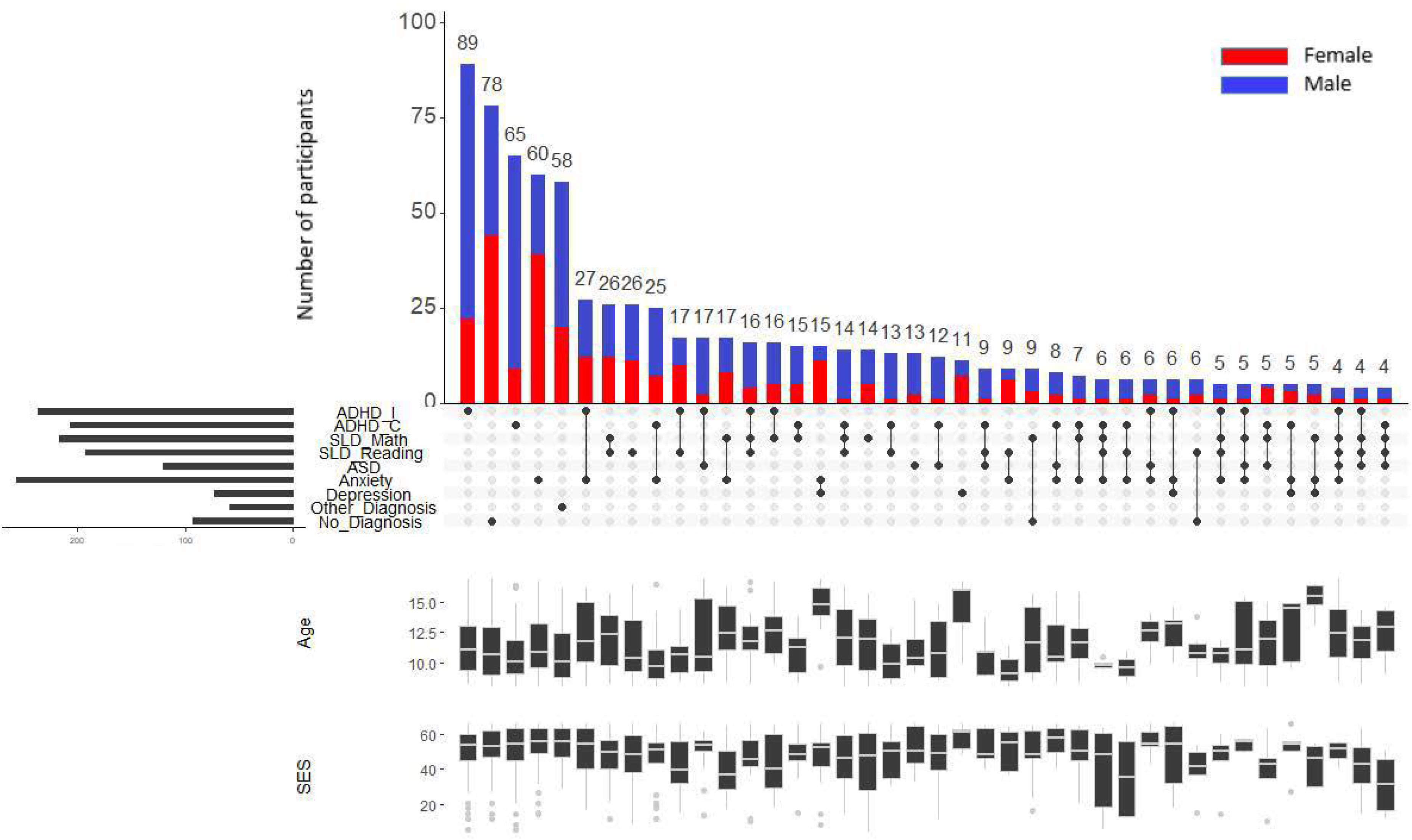

### Inter-measure Reliability

Single-fixed raters intraclass correlation coefficients (ICC) were higher between PC1 and each of the four individual PS measures than between the four individual PS measures (See Table 2). WISC-SS Clinician had highest reliability with WISC-Coding, followed by WISC-SS EEG, and had the lowest reliability with NIH-PC. WISC-Coding had moderate reliability with WISC-SS EEG, and low reliability with NIH-PC. NIH PC also had low reliability with WISC-SS EEG. Given the low to moderate reliability between measures (ranging between .14 and .66), and the strong reliability between PC1 and all four measures, PC1 was used as the main PS construct for all analyses.

**Table 2:**
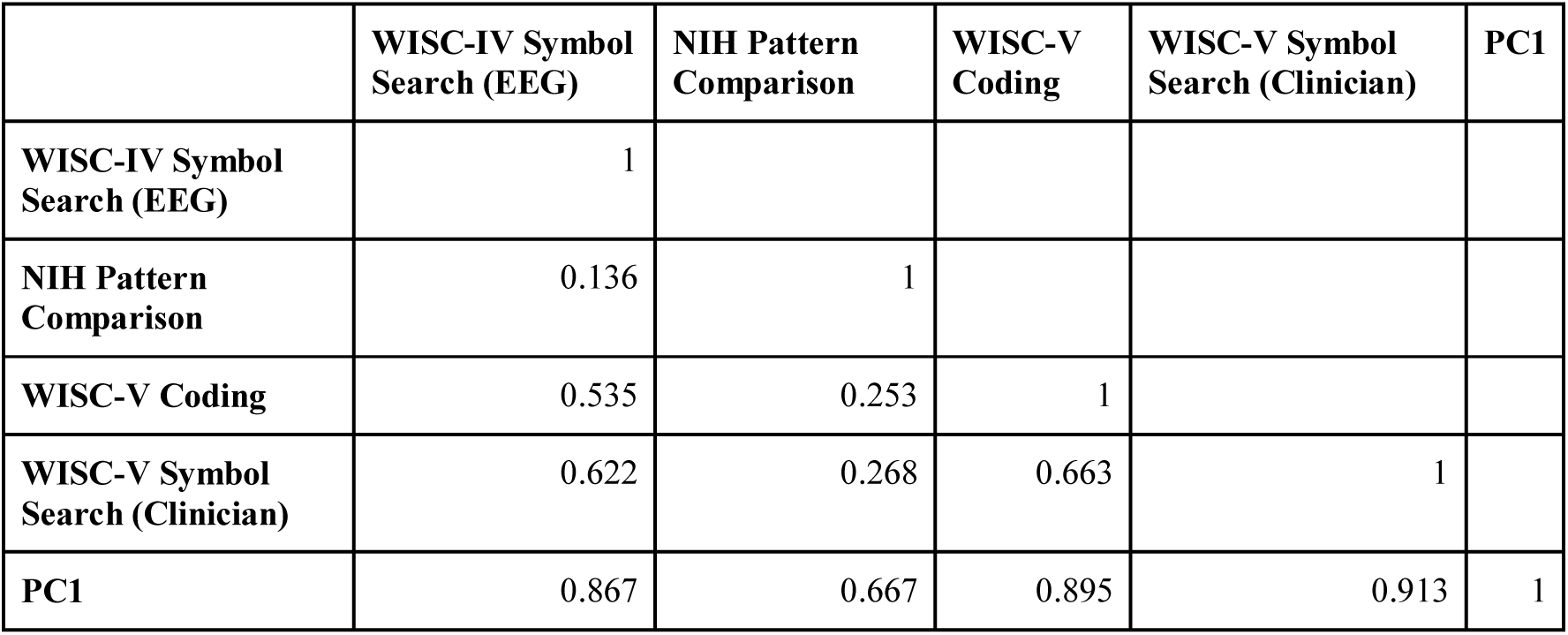
Reliabilities calculated between the four different processing speed tasks, as well as the principal component-based composite score (i.e., first principal component - PC1).

The first principal component (PC1) accounted for 59.16% of the total variance and is included in the primary analysis. All four individual PS measures loaded positively onto PC1, with NIH-PC having the weakest relationship. Pearson correlations between PC1 and the four measures indicated strong associations: WISC-SS Clinician r(841) = 0.84, p < .001; WISC-Coding r(841) = 0.81, p < .001; WISC-SS EEG r(841) = 0.79, p < .001; NIH-PC r(841) = 0.60, p < .001).

The second principal component (PC2) accounted for 19.07% of the variance, which was less than 25%, i.e. the percentage of variance explained if all four principal components accounted for the same percentage of variance. For this reason, PC2 was not included in primary analyses (see supplemental materials for analyses of how the tasks load onto PC1 and PC2, as well as assessment of PC2 and its associations with cognitive factors and psychopathology).

### Accounting for Potential Confounds: Age, Sex, SES, Motor Control, Full Scale IQ

The false discovery rate-corrected Pearson correlations between parent and self-reported questionnaires of psychopathology and PC1 resulted in positive correlations between PC1 and sex (r(841) = 0.20, p = <0.001) and SES as measured by the Barratt (r(836) = 0.08, p = <0.001). PC1 and age were not significantly correlated (r(841) = -0.06, p = 0.10), likely due to the fact that the four subtests are standardized scores that take age into account already. Collection site was also not significantly associated with PC1 (r(841) = 0.019, p = 0.5731). The correlation between PC1 and motor control, as measured by Grooved Pegboard task using the participant’s dominant hand, was significant (r(837) = 0.38, p = <.001), suggesting motor control might be a possible confound. Thus, we included motor control, together with age, sex, SES, and collection site as covariates in multiple regression analyses.

A common question in the processing speed literature is whether it is necessary to account for the contributions of other components of intelligence to processing speed related findings. To accomplish this, some studies control for full-scale IQ when testing for associations with PS, while others view this as too conservative, as processing speed is one of its determinants. We considered the possibility of using the general ability index (GAI), which is similar to FSIQ but does not include processing speed or working memory in its calculation; this is commonly used clinically to provide a potentially more independent depiction of general intelligence in individuals with PS deficits. However, the correlation between WISC-GAI and WISC-FSIQ was extremely high (r(818) = 0.96, p < .001), indicating that GAI does not adequately separate general intelligence from PS. For this reason, as well as keeping consistent with other studies in the field, WISC-FSIQ was used as the measure of general intelligence for all analyses. The correlation between WISC-FSIQ and PC1 was strong (r(841) = 0.61, p = <.001).

Note that the primary results reported in the following sections of the manuscript are not corrected for FSIQ due to concerns about potential overcorrection. However, we do report in each section whether or not correction for FSIQ modified our findings and provide more comprehensive reporting in the supplementary materials.

### Associations with Psychopathology: Categorical Diagnoses

*Categorical.* Multiple linear regression analyses were performed to test for specific associations between processing speed (PC1) and the DSM-V diagnoses assessed using the KSADS. Significant negative relationships were detected for the presence of each, ADHD-Inattentive type, ADHD-Combined type, and both SLD subgroups, after accounting for age, sex, SES, motor skills, and collection site (F= 27.56, df = 786, adjusted R^2^ = .29; Table 3).

**Table 3:** Processing speed regressed on categorical diagnostic labels—comparing when the 7 disorders are included in one regression model predicting processing speed (as indexed by PC1), and when each one is modeled alone. For all regression models, demographic variables (age, sex, SES, collection site) and grooved pegboard were included as nuisance covariates.

|  | Each Diagnosis Modeled in Separate Models |  |  |  |  |  | *Diagnoses Modeled Together* |  |  |
| --- | --- | --- | --- | --- | --- | --- | --- | --- | --- |
|  | *beta* | *SE* | *p-value* | *F statistic* | *df* | *Adjusted R<sup>2</sup>* | *beta* | *SE* | *p-value* |
| ADHD-I | -0.444 | 0.106 | 3.09E-05*** | 32.34054 | 792 | 0.19 | -0.546 | 0.109 | 6.32E-07*** |
| ADHD-C | -0.175 | 0.115 | 0.13 | 29.24027 | 792 | 0.18 | -0.322 | 0.118 | 0.0065** |
| ASD | -0.316 | 0.140 | 0.025* | 29.79968 | 792 | 0.18 | -0.153 | 0.133 | 0.25 |
| SLD-Math | -0.981 | 0.109 | 1.86E-18*** | 45.16474 | 792 | 0.25 | -0.844 | 0.111 | 6.87E-14*** |
| SLD-Reading | -0.674 | 0.116 | 9.13E-09*** | 35.62329 | 792 | 0.21 | -0.444 | 0.115 | 0.00011*** |
| Anxiety | 0.048 | 0.106 | 0.65 | 28.81604 | 792 | 0.17 | 0.0232 | 0.101 | 0.82 |
| Depression | 0.172 | 0.180 | 0.34 | 28.95878 | 792 | 0.17 | 0.144 | 0.171 | 0.4 |

Sex significantly predicted PC1 (b = 0.414, SE = 0.097, p = 2.3E-05***) indicating that females are faster at PS tasks. SES (b = 0.009, SE= 0.003, p = 0.006**), collection site (b = 0.23, SE = 0.10, p = 0.017*) and motor control (b = 0.346, SE = 0.037, p = 1.56E-19***) also significantly contribute to PS outcomes in the categorical model. Age was not a significant predictor of PC1 (b = -0.014, SE = 0.020, p = 0.483), likely due to PC1 being age corrected through standardized scores. Accounting for FSIQ does not change significance in diagnosis predicting PS deficit, with the exception of SLD-reading subgroup becoming nonsignificant.

Included in Table 3 are analyses that show which diagnoses significantly predict PC1 when that sole diagnosis, along with the demographic and motor variables, is entered into a linear regression predicting PC1.

### Associations with Symptomatology: Dimensional Measures

Correlation analyses (corrected for multiple comparisons using false discovery rate) showed that PC1 exhibited significant negative correlations with symptoms of inattention (r(839 = - 0.21, p < 0.001) and hyperactivity (r(839) = -0.13, p < 0.001) as measured by the SWAN subscales. In addition, the three measures of ASD were negatively correlated with PC1 (ASSQ r(839) = -0.21, p < 0.001; SCQ r(841) = -0.18, p < 0.001; SRS r(841) = -0.21, p < 0.001).

These measures were then entered into a linear regression analysis to analyze relationships among these variables; age, sex, SES, motor control and collection site were included as confound variables (F = 21.27, df = 783, adjusted R^2^ = .20; Table 4). The SWAN Inattention subscale (SWAN-IN) was found to have as strong, independent negative association with PC1, as well as the ASSQ. In addition, sex, SES, collection site, and motor control were significant. When FSIQ is included in the model, the significance and direction of the results remain the same. If the questionnaires are examined in isolation, with each of the five subscales predicting PC1 while accounting for demographic variables in a linear regression, four of the five scales significantly predict PC1, with SCQ being borderline (Table 4).

**Table 4:** Processing speed regressed on dimensional measures — comparing when the 5 questionnaires are included in one regression model predicting processing speed (as indexed by PC1), and when each one is modeled alone. For all regression models, demographic variables and grooved pegboard were included as nuisance covariates.

|  | Each Questionnaire Modeled in Separate Model |  |  |  |  |  | *Questionnaires Modeled Together* |  |  |
| --- | --- | --- | --- | --- | --- | --- | --- | --- | --- |
|  | *beta* | *SE* | *p-value* | *F statistic* | *df* | *Adjusted R<sup>2</sup>* | *beta* | *SE* | *p-value* |
| SWAN-IN | -0.221 | 0.04 | 6.04E-08*** | 34.3199 | 787 | 0.18 | -0.22 | 0.05 | 1.71E-05*** |
| SWAN-HY | -0.13 | 0.04 | 0.0026** | 30.10752 | 787 | 0.2 | 0.026 | 0.05 | 0.64 |
| ASSQ | -0.02 | 0.005 | 0.00026*** | 30.99247 | 787 | 0.18 | -0.019 | 0.009 | 0.025* |
| SRS | -0.005 | 0.002 | 0.00086*** | 30.52574 | 787 | 0.18 | 0.0023 | 0.003 | 0.45 |
| SCQ | -0.019 | 0.01 | 0.0503 | 29.03677 | 787 | 0.18 | 0.0025 | 0.013 | 0.85 |

### Post-Hoc Examination of Associations with Inattention

Given the strong relationship between the SWAN-Inattention subscale and PC1, we included the SWAN-Inattentive subscale score as a predictor in the categorical regression model examining clinician diagnoses as predictors of PC1 (F = 25.84, df = 783, adjusted R^2^ = 0.29). When inattentive symptoms were included in the model, the relationship between ADHD-C and PC1 was no longer significant (p = 0.12). This indicates that the relationship between ADHD-C and PS deficit is reliant upon inattention, as opposed to other executive function (EF) deficit. ADHD-I continued to significantly predict PC1 (p = 0.0005). The two SLD subgroups also continued to significantly predict PC1 after accounting for inattentive symptoms (SLD-Reading p = 0.0002; SLD-Math p = 3.34E-13). This indicates that there are factors that account for the relationship between SLD and PS deficit apart from inattentive symptoms. When FSIQ is included in the model, both subtypes of ADHD predict PC1, and only the SLD-Math subgroup significantly predicts PC1.

## DISCUSSION

The present study leveraged the transdiagnostic Healthy Brain Network Biobank to explore the relationship between PS deficits and psychopathology, both with respect to DSM-V diagnoses and commonly used dimensional psychiatric assessments. Consistent with prior work, our examination of individual associations between PS and each, DSM-V disorders and their related dimensional measures, implicated PS impairments in attention deficit/hyperactivity (ADHD)^7, 8, 13, 29, 81, 82^, autism spectrum (ASD)^2, 7, 8, 33, 44, 46^ and specific learning disabilities (SLD)^7, 9, 12, 14, 15, 41^. However, simultaneous analyses suggested more specific associations with ADHD and learning disorders. Among individuals with ADHD, associations appeared to be most related to symptoms of inattention, as dimensional measures of hyperactivity did not exhibit specific significant association with PS, and relations with ADHD-C diagnosis do not remain if dimensional measures of inattention are considered. Finally it is worth noting that consistent with prior work^82–85^, being male was an independent negative predictor for processing speed, while higher socioeconomic status was a positive predictor.

In contrast to subsets of reports in the emerging literature for PS associations with psychopathology^2, 8, 44, 46, 56, 57, 59^, we did not find any specific associations between PS and the diagnoses of ASD, anxiety, and depression. Furthermore, only ASD exhibited a significant association with PS prior to accounting for other diagnoses. While this may reflect differences in sampling strategies between the present work and other studies, it is also possible that these findings could be explained by comorbid diagnosis of ADHD or the presence of inattentive symptoms^53, 86, 87^. The same pattern emerged dimensionally, as the ASD questionnaires significantly predicted PS deficit when examined in isolation, but largely failed to do so when inattentive symptoms were included in the model. At a minimum, our findings suggest the importance of future efforts to consider transdiagnostic perspectives when attempting to establish associations - particularly for diagnoses known to have a high frequency of co-occurring ADHD (e.g., ASD, anxiety disorders) and the advantages of working within a multiple deficit framework.

As largely indicated in the literature^9–11, 23, 25^, the presence of a SLD significantly predicted PS deficit. Reading and mathematics impairment subgroups were each related to PS deficit when examined through a categorical lens. Interestingly, both subtypes predicted PS even after controlling for inattention, suggesting that other factors could mediate the relationship between SLD and PS deficit. Also within the categorical framework, the SLD-Reading subgroup failed to predict PS after controlling for FSIQ. This could be due to the strong relationship between intelligence and learning; it is also worth keeping in mind that there are speeded components of FSIQ, including the PS tasks, Visual Puzzles, Figure Weights and Block Design. These collective findings support previous research showing a relationship between MD, RD, and PS deficit^21, 22, 24, 25^. A wider array of PS tasks would be necessary to more fully unpack the relationships between SLD, inattention, and domain specific PS difficulties.

The prominence of findings for ADHD in the present work is not surprising, though relations with PS were found to be notably more consistent across analyses for ADHD-I than ADHD-C. Recent reports have associated ADHD-related deficits in PS to a number of other areas of impairment in ADHD, including externalizing behaviors^31, 35, 42, 88^, inhibition deficits^35, 79, 81, 89– 91^, and working memory deficits^1, 91, 92^. This study points to inattention as having the driving role in the relationships between ADHD subtypes and PS deficits, supporting previous research highlighting inattention^34–36, 38, 41^. In the categorical model, ADHD-C predicts PS deficits, but this relationship becomes non-significant when accounting for inattentive symptoms. Similarly, in the dimensional analysis, hyperactive symptoms did not significantly predict PC1 when inattention and autistic symptoms were included. Given that the role of ADHD-C in PS deficits is highly debated, and that the nature of how hyperactive symptoms and inattentive symptoms interact with each other and with PS deficits is a complex topic, further research into these specific symptoms and their interaction with each other and with PS is needed.

In line with previous research^83–85, 93^, females generally had higher PS scores than males, and sex was a significant predictor of PS in all analyses. This was found to be true even after accounting for all dimensional measures of psychopathology. Similarly, motor control was strongly and frequently associated with PS, which underlines the importance of accounting for motor deficits, as these are common in children with SLD, ADHD, and ASD^34, 46, 54, 60, 79, 90^. In accordance with previous work on the relationship between SES and PS^94–96^, SES was also significantly positively associated with PS throughout all analyses. There were no significant associations with age, which is typically seen as being very important in PS development, though this is likely due to the use of standardized PS scores, which are age-normalized.

For the purposes of the present work, we made use of a single construct derived from scores obtained using all four processing speed tasks included in the Healthy Brain Network to have the most robust measure of processing speed. However, an important caution that emerges from the present work is the low to moderate reliability among the different processing speed tasks. Most notable, were those between the NIH Toolbox Pattern Comparison task and the WISC-based instruments (i.e., Symbol Search, Coding), which ranged between 0.136 and 0.268. These are in a similar range to prior work^97–99^, though somewhat lower - likely reflecting the notably larger sample size in the present work, as well as possible increases in within-individual variation that would be expected with the presence of mental health and learning disorders in our sample. The Pattern Comparison task is unique in that it does not have the extent of working memory requirements or motor demands present in Symbol Search or Coding, likely explaining the larger disparity in reliabilities with other tasks. Regardless of which task one employs, our findings regarding the reliability among processing speed tasks raises cautions for the study of these tasks in small samples, as the lower reliabilities will inherently increase false positive and negative findings. The supplemental materials indicate overall consistency with PC1, though a few variances that should be considered when interpreting task results.

The present study has several limitations that should be considered when evaluating its clinical significance and replicability. The first is that this study uses a convenience sample, created from a dataset that already existed and was not designed specifically for this analysis. Therefore, there is no randomization of task order. Secondly, the PC1 variable accounted for 60% of the variance in the four tasks; the ideal variance for a principal component is 70-80%. This relatively low variance is due to an uneven capture of the four tasks, with NIH Toolbox having a weaker relationship and less in common with the other three tasks. Third, PC1 is not a set construct-- the value of this construct depends on both the individual performance on the four PS measures, as well as the performance of the rest of the sample used to calculate it. Similarly, a fourth limitation is the necessity for administration of all four tasks in order for these results to generalize to clinical application. Finally, there were confounding environmental factors during the WISC-IV computerized Symbol Search task, as participants sat in a dark room with a wet EEG cap on their heads.

Looking forward, future research would benefit from further exploration of the relationship between PS deficits and specific neuropsychological deficits underlying ADHD and SLD symptomatology. Given the prevalence of PS deficits across a range of disorders in the present work and the larger clinical literature, there may be a value to increased consideration of processing speed as a specific construct in transdiagnostic research frameworks (e.g., at the present time, the Research Domain Criteria (RDoC) project does not currently include PS in the cognitive systems domain). Additionally, from a clinical perspective, the findings of the present work motivate an increased focus on the screening of processing speed impairments in children with mental health disorders, particularly those exhibiting impairments in attention.

## Acknowledgements

The work presented here was primarily supported by gifts to the Child Mind Institute from Phyllis Green, Randolph Cowen, and Joseph Healey, as well as NIMH awards to Dr. Milham (U01MH099059, R01MH091864). Additionally, we would like to thank Aki Nikolaidis and Ting Xu for their advice regarding our analytic strategy, as well as Nicolas Langer for his advice in thinking through our study. We would like to thank and acknowledge the thousands of families who participated in this ongoing project and the generous philanthropic supports who made it possible, as well as the mental health organizations, service providers, and clinicians across NYC who continue to refer families to our team.

