## Supplemental Materials for "Associations between Processing Speed and Psychopathology in a Transdiagnostic, Pediatric Sample"

Supplemental Material

### Supplementary Text

**S1:** This table shows the variable loadings of the four subtests onto the four PCA components.

**S2-4:** Multiple linear regressions examined the same analyses run in the main paper on PC1, while also including FSIQ as a covariate. Though we believe results should be examined without FSIQ given the intricate relationship between intelligence and many of these variables, these tables are included to show that the major findings largely remained intact when overall intelligence was taken into account. Table S2 examines how categorical diagnosis from HBN clinicians relates with PC1. Table S3 examines which dimensional symptoms from parent and self-report questionnaires are related to PC1. Table S4 shows results from the regression analysis which examines relationships between categorical diagnoses and PC1 while controlling for inattentive symptoms.

**S5-8:** The four tables in this section outline the same analyses run on PC1, but with mental health symptoms and diagnoses predicting the four processing speed tasks individually. These tables assist with comparing results across the literature for publications that only use one of the PS tests. Table S5 shows the relationships between the four PS tests and variables of interest, including confounding variables and dimensional symptoms. The following three tables (S6-8) show the results of multiple linear regression models predicting the four PS tests individually, with and without FSIQ as an added covariate.

**S9-11:** S9-11 tables show the relationship between PC2 and pediatric mental health. PC2 explained only 19% of the variance of the four tasks and thus was not included in the main manuscript. Table S9 shows correlations between PC2, confounding variables and dimensional symptoms. Table S10 shows the linear regression examining relationships between categorical diagnosis and PC2, with and without including FSIQ as an added covariate. Finally, Table S11 shows the linear regression examining how dimensional symptoms relate with PC2, with and without FSIQ as an added covariate. Given that PC2 is not related to inattentive symptoms, further regressions specifically examining inattention were not included.

For all analyses, please refer to the following to indicate level of significance: p < 0.05 *, p < 0.01**, p < 0.001***

| **PCA Variable Loadings** | | | | |
| --- | --- | --- | --- | --- |
|  | **PC1** | **PC2** | **PC3** | **PC4** |
| **WISC-Coding** | 0.5336653 | -0.1405348 | 0.6610039 | -0.50845374 |
| **WISC-SS** | 0.5497116 | -0.1753970 | 0.1380940 | 0.80497399 |
| **NIH PC** | 0.3905519 | 0.9063796 | -0.1553503 | -0.04256243 |
| **WISC-SS EEG** | 0.5103801 | -0.3577176 | -0.7210202 | -0.30278723 |

### **Table S1**: Variable loadings for the four subtests onto the PCA components

### **Table S2**: PC1 regressed on categorical diagnostic labels, with FSIQ included as a covariate

F statistic = 39.82, df =785, adjusted R^2^ = 0.39

| variable | b value | SE | p value |
| --- | --- | --- | --- |
| (Intercept) | -3.591 | 0.4265 | 1.78E-16*** |
| ADHD-I | -0.4422 | 0.1011 | 1.39E-05*** |
| ADHD-C | -0.2837 | 0.1092 | 0.009556** |
| ASD | -0.1383 | 0.1232 | 0.2621 |
| SLD-Math | -0.3774 | 0.1102 | 0.0006513*** |
| SLD-Reading | -0.08461 | 0.1105 | 0.4442 |
| Anxiety | -0.01889 | 0.09313 | 0.8393 |
| Depression | 0.06272 | 0.1581 | 0.6917 |
| Sex | 0.4091 | 0.08994 | 6.26E-06*** |
| Age | -0.01238 | 0.01861 | 0.5061 |
| Site | 0.04009 | 0.0901 | 0.6565 |
| SES | 9.02E-05 | 0.003048 | 0.9764 |
| Motor Control | 0.1933 | 0.03692 | 2.12E-07*** |
| WISC-FSIQ | 0.04095 | 0.003566 | 2.59E-28*** |

### **Table S3:** PC1 regressed on dimensional measures, with FSIQ included as a covariate

F statistic = 44.57, df = 782, adjusted R^2^ = 0.38

| variable | b value | SE | p value |
| --- | --- | --- | --- |
| (Intercept) | -4.38 | 0.3911 | 4.18E-27*** |
| SWAN-IN | -0.09857 | 0.04616 | 0.03302* |
| SWAN-HY | 0.01423 | 0.04851 | 0.7693 |
| ASSQ | -0.02532 | 0.007635 | 0.0009552*** |
| SRS | 0.001578 | 0.002678 | 0.5558 |
| SCQ | 0.01771 | 0.01181 | 0.134 |
| Sex | 0.4312 | 0.08984 | 1.90E-06*** |
| Age | -0.01977 | 0.01831 | 0.2806 |
| SES | -0.000572 | 0.003119 | 0.8545 |
| Site | 0.01253 | 0.08913 | 0.8882 |
| Motor Control | 0.1836 | 0.0367 | 6.99E-07*** |
| WISC-FSIQ | 0.04717 | 0.003191 | 8.60E-44*** |

### **Table S4:** PC1 regressed on categorical diagnostic labels with SWAN-inattention and FSIQ included as covariates

F statistic = 36.93, df = 782, adjusted R^2^ = 0.39

| variable | b value | SE | p value |
| --- | --- | --- | --- |
| (Intercept) | -3.582 | 0.4323 | 5.14E-16*** |
| ADHD-I | -0.4059 | 0.1132 | 0.000356*** |
| ADHD-C | -0.2473 | 0.1213 | 0.04179* |
| ASD | -0.1318 | 0.1236 | 0.2867 |
| SLD-Math | -0.3821 | 0.1107 | 0.0005829*** |
| SLD-Reading | -0.07789 | 0.1107 | 0.4819 |
| Anxiety | -0.01738 | 0.09335 | 0.8523 |
| Depression | 0.07721 | 0.159 | 0.6274 |
| Sex | 0.3999 | 0.09021 | 1.06E-05*** |
| Age | -0.01249 | 0.01867 | 0.5035 |
| Site | 0.04107 | 0.0902 | 0.649 |
| SES | 0.0004573 | 0.003067 | 0.8815 |
| Motor Control | 0.1932 | 0.03695 | 2.20E-07*** |
| SWAN-IN | -0.02529 | 0.04252 | 0.5522 |
| WISC-FSIQ | 0.04063 | 0.003605 | 2.08E-27*** |

### **Table S5**: Correlations among processing speed tasks, potential confounds and dimensional symptoms

| **Variable** | **WISC-Coding** | **WISC-SS Clinician** | **WISC-SS EEG** | **NIH PC** |
| --- | --- | --- | --- | --- |
| Age | r(1458) = -0.126, *** | r(1450) = -0.086, ** | r(841) = -0.127, *** | r(841) = 0.159, *** |
| Sex | r(1458) = 0.160, *** | r(1450) = 0.068, * | r(841) = 0.095, ** | r(841) = 0.074, * |
| SES | r(1331) = 0.147, *** | r(1319) = 0.151, *** | r(836) = 0.152, *** | r(836) = 0.070 |
| Motor Control | r(1450) = 0.324, *** | r(1437) = 0.321, *** | r(837) = 0.293, *** | r(837) = 0.201, *** |
| FSIQ | r(1474) = 0.583, *** | r(1462) = 0.536, *** | r(844) = 0.543, *** | r(844) = 0.251, *** |
| SWAN-IN | r(1341) = -0.172, *** | r(1329) = -0.162, *** | r(840) = -0.200, *** | r(840) = -0.090, * |
| SWAN-HY | r(1341) = -0.105, *** | r(1329) =-0.088, ** | r(840) = -0.133, *** | r(840) = -0.045 |
| ASSQ | r(1342) = -0.139, *** | r(1330) = -0.142, *** | r(840) = -0.182, *** | r(840) = -0.069 |
| SRS | r(1355) = -0.150, *** | r(1343) = -0.165,*** | r(842) = -0.189, *** | r(842) = -0.072 |
| SCQ | r(1353) = -0.137, *** | r(1341) = -0.143, *** | r(842) = -0.176, *** | r(842) = 0.002 |

*all results corrected for multiple comparisons using false discovery rate

Note: though only these five scales are included above, in keeping with consistency with the main paper, values were also examined for the rest of the dimensional symptoms (MFQ, SCARED, ARI). There were no additional associations, apart from SCARED-Parent and ARI-Self being correlated with WISC-SS EEG, both at the * level.

### **Table S6:** Each of the four processing speed tasks regressed separately on categorical diagnostic labels

**Without FSIQ:**

| variable | **WISC-Coding** | **WISC-SS Clinician** | **WISC-SS EEG** | **NIH PC** |
| --- | --- | --- | --- | --- |
| (Intercept) | 9.34E-151*** | 4.27E-141*** | 2.79E-155*** | 6.54E-47*** |
| ADHD-I | 2.78E-07*** | 0.0053** | 0.0001*** | 0.007** |
| ADHD-C | 0.034* | 0.12 | 0.027* | 0.045* |
| ASD | 0.06 | 0.23 | 0.23 | 0.20 |
| SLD-Math | 1.11E-13*** | 2.40E-08*** | 4.17E-11*** | 0.13 |
| SLD-Reading | 0.00013*** | 0.0069** | 0.0014** | 0.23 |
| Anxiety | 0.569 | 0.536 | 0.51 | 0.91 |
| Depression | 0.47 | 0.70 | 0.66 | 0.34 |
| Sex | 4.70E-12*** | 0.002** | 0.30 | 0.44 |
| Age | 0.06 | 0.25 | 0.0097** | 1.25E-05*** |
| Site | 0.94 | 0.036* | 0.00017*** | 0.42 |
| SES | 0.047* | 0.0014** | 0.1315 | 0.34 |
| Motor Control | 1.13E-10*** | 1.05E-14*** | 4.75E-11*** | 8.52E-08*** |
| F statistic | 26.16 | 17.18 | 16.89 | 6.75 |
| df | 786 | 786 | 786 | 786 |
| Adjusted R^2^ | 0.27 | 0.20 | 0.19 | 0.08 |

**With FSIQ:**

| variable | **WISC-Coding** | **WISC-SS Clinician** | **WISC-SS EEG** | **NIH PC** |
| --- | --- | --- | --- | --- |
| (Intercept) | 3.17E-41*** | 4.27E-42*** | 4.47E-44*** | 5.51E-11*** |
| ADHD-I | 5.36E-06*** | 0.029* | 0.0021** | 0.019* |
| ADHD-C | 0.051 | 0.17 | 0.041* | 0.057 |
| ASD | 0.061 | 0.25 | 0.25 | 0.18 |
| SLD-Math | 0.00017*** | 0.010* | 0.003** | 0.95 |
| SLD-Reading | 0.28 | 0.62 | 0.63 | 0.97 |
| Anxiety | 0.83 | 0.34 | 0.76 | 0.97 |
| Depression | 0.75 | 0.96 | 0.97 | 0.43 |
| Sex | 3.48E-13*** | 0.0015** | 0.29 | 0.45 |
| Age | 0.057 | 0.26 | 0.0078** | 8.93E-06*** |
| Site | 0.081 | 0.48 | 0.033* | 0.95 |
| SES | 0.67 | 0.19 | 0.38 | 0.96 |
| Motor Control | 0.004** | 1.55E-06*** | 0.0023** | 0.0003*** |
| WISC-FSIQ | 4.23E-22*** | 2.52E-14*** | 1.33E-21*** | 6.91E-05*** |
| F statistic | 34.80 | 21.70 | 24.94 | 7.58 |
| df | 785 | 785 | 785 | 785 |
| Adjusted R^2^ | 0.36 | 0.25 | 0.28 | 0.10 |

### **Table S7:** Each of the four processing speed tasks regressed separately on dimensional measures

**Without FSIQ:**

| variable | **WISC-Coding** | **WISC-SS Clinician** | **WISC-SS EEG** | **NIH PC** |
| --- | --- | --- | --- | --- |
| (Intercept) | 9.54E-129*** | 9.57E-128*** | 8.18E-141*** | 2.61E-39*** |
| SWAN-IN | 0.0002*** | 0.0033** | 3.93E-05*** | 0.095 |
| SWAN-HY | 0.65 | 0.86 | 0.98 | 0.40 |
| ASSQ | 0.061 | 0.018* | 0.14 | 0.50 |
| SRS | 0.61 | 0.21 | 0.54 | 0.71 |
| SCQ | 0.99 | 0.53 | 0.84 | 0.086 |
| Sex | 4.11E-12*** | 0.0064** | 0.36 | 0.19 |
| Age | 0.024* | 0.17 | 0.004** | 1.98E-06*** |
| SES | 0.006** | 0.00018*** | 0.044* | 0.16 |
| Site | 0.79 | 0.08 | 0.0005*** | 0.44 |
| Motor Control | 1.90E-11*** | 6.67E-16*** | 3.80E-12*** | 1.05E-08*** |
| F statistic | 18.64 | 15.43 | 13.20 | 7.30 |
| df | 783 | 783 | 783 | 783 |
| Adjusted R^2^ | 0.18 | 0.15 | 0.13 | 0.07 |

**With FSIQ:**

| variable | **WISC-Coding** | **WISC-SS Clinician** | **WISC-SS EEG** | **NIH PC** |
| --- | --- | --- | --- | --- |
| (Intercept) | 4.98E-36*** | 2.67E-42*** | 1.31E-45*** | 2.20E-10*** |
| SWAN-IN | 0.10 | 0.21 | 0.03* | 0.42 |
| SWAN-HY | 0.78 | 0.98 | 0.86 | 0.44 |
| ASSQ | 0.006** | 0.0024** | 0.025* | 0.35 |
| SRS | 0.74 | 0.25 | 0.66 | 0.64 |
| SCQ | 0.24 | 0.83 | 0.39 | 0.032* |
| Sex | 1.44E-14*** | 0.003** | 0.30 | 0.18 |
| Age | 0.012* | 0.14 | 0.0016** | 1.49E-06*** |
| SES | 0.51 | 0.26 | 0.23 | 0.91 |
| Site | 0.031* | 0.68 | 0.042* | 0.91 |
| Motor Control | 0.016* | 3.10E-06*** | 0.0035** | 0.0001*** |
| WISC-FSIQ | 5.02E-37*** | 1.05E-22*** | 3.78E-32*** | 2.45E-06*** |
| F statistic | 37.15 | 25.15 | 28.20 | 8.87 |
| df | 782 | 782 | 782 | 782 |
| Adjusted R^2^ | 0.34 | 0.26 | 0.28 | 0.11 |

### **Table S8:** Each of the four processing speed tasks regressed separately on categorical diagnostic labels, including SWAN-inattention as a covariate

**Without FSIQ:**

| variable | **WISC-Coding** | **WISC-SS Clinician** | **WISC-SS EEG** | **NIH PC** |
| --- | --- | --- | --- | --- |
| (Intercept) | 1.32E-149*** | 7.03E-140*** | 2.69E-155*** | 4.49E-46*** |
| ADHD-I | 6.39E-05*** | 0.08 | 0.040* | 0.022* |
| ADHD-C | 0.18 | 0.48 | 0.44 | 0.083 |
| ASD | 0.079 | 0.29 | 0.33 | 0.20 |
| SLD-Math | 4.63E-13*** | 5.05E-08*** | 3.14E-10*** | 0.13 |
| SLD-Reading | 0.0002*** | 0.0099** | 0.0021** | 0.25 |
| Anxiety | 0.57 | 0.54 | 0.57 | 0.87 |
| Depression | 0.40 | 0.58 | 0.47 | 0.33 |
| Sex | 9.13E-12*** | 0.0033** | 0.41 | 0.46 |
| Age | 0.053 | 0.23 | 0.0076** | 1.45E-05*** |
| Site | 0.98 | 0.04* | 0.0002*** | 0.43 |
| SES | 0.056 | 0.0013** | 0.17 | 0.30 |
| Motor Control | 1.93E-10*** | 2.03E-14*** | 9.26E-11*** | 1.04E-07*** |
| SWAN-IN | 0.20 | 0.13 | 0.0061** | 0.89 |
| F statistic | 24.17 | 16.12 | 16.25 | 6.23 |
| df | 783 | 783 | 783 | 783 |
| Adjusted R^2^ | 0.27 | 0.20 | 0.20 | 0.08 |

**With FSIQ:**

| variable | **WISC-Coding** | **WISC-SS Clinician** | **WISC-SS EEG** | **NIH PC** |
| --- | --- | --- | --- | --- |
| (Intercept) | 4.88E-40*** | 4.06E-41*** | 4.09E-44*** | 2.04E-10*** |
| ADHD-I | 4.59E-05*** | 0.094 | 0.044* | 0.02* |
| ADHD-C | 0.077 | 0.32 | 0.26 | 0.060 |
| ASD | 0.062 | 0.27 | 0.30 | 0.19 |
| SLD-Math | 0.0002*** | 0.009** | 0.0037** | 0.99 |
| SLD-Reading | 0.29 | 0.67 | 0.65 | 0.99 |
| Anxiety | 0.82 | 0.36 | 0.80 | 0.99 |
| Depression | 0.75 | 0.89 | 0.83 | 0.45 |
| Sex | 4.63E-13*** | 0.0022** | 0.37 | 0.45 |
| Age | 0.057 | 0.26 | 0.007** | 9.12E-06*** |
| Site | 0.080 | 0.47 | 0.030* | 0.96 |
| SES | 0.70 | 0.15 | 0.40 | 0.96 |
| Motor Control | 0.004** | 1.62E-06*** | 0.0020** | 0.0003*** |
| SWAN-IN | 0.96 | 0.62 | 0.12 | 0.67 |
| WISC-FSIQ | 1.10E-21*** | 7.63E-14*** | 2.51E-20*** | 6.11E-05*** |
| F statistic | 32.14 | 20.20 | 23.25 | 7.06 |
| df | 782 | 782 | 782 | 782 |
| Adjusted R^2^ | 0.35 | 0.25 | 0.28 | 0.10 |

### **Table S9:** Correlations between PC2 and four individual PS measures, potential confounds, and dimensional symptoms

| **Variable** | **R value** | **P value** | **df** |
| --- | --- | --- | --- |
| WISC-Coding | -0.123 | 0.0003*** | 844 |
| WISC-SS Clinician | -0.153 | 7.61E-06*** | 844 |
| WISC-SS EEG | -0.312 | 1.30E-20*** | 844 |
| NIH PC | 0.792 | 8.44E-183*** | 844 |
| Age | 0.248 | 1.04E-12*** | 841 |
| Sex | -0.037 | 0.364 | 841 |
| SES | 0.152 | 0.224 | 836 |
| Motor Control | -0.031 | 0.460 | 837 |
| FSIQ | -0.147 | 1.67E-05*** | 844 |
| SWAN-IN | 0.048 | 0.226 | 840 |
| SWAN-HY | 0.045 | 0.259 | 840 |
| ASSQ | 0.072 | 0.053 | 840 |
| SRS | 0.068 | 0.069 | 842 |
| SCQ | 0.136 | 0.0001*** | 842 |

*all results corrected for multiple comparisons using false discovery rate

### **Table S10:** PC2 regressed on categorical diagnostic labels

**Without FSIQ:** F statistic = 5.98, DF = 786, adjusted R^2^ = 0.07

| variable | b value | SE | p value |
| --- | --- | --- | --- |
| (Intercept) | -0.9278 | 0.1942 | 2.11E-06*** |
| ADHD-I | -0.01212 | 0.07297 | 0.8681 |
| ADHD-C | -0.05369 | 0.07909 | 0.4975 |
| ASD | 0.2002 | 0.08929 | 0.02525* |
| SLD-Reading | 0.2217 | 0.07425 | 0.002923** |
| SLD-Math | 0.07385 | 0.0768 | 0.3365 |
| Anxiety | -0.005792 | 0.06743 | 0.9316 |
| Depression | 0.07433 | 0.1145 | 0.5163 |
| Sex | -0.07299 | 0.06517 | 0.2631 |
| Age | 0.08039 | 0.01348 | 3.75E-09*** |
| SES | -0.06147 | 0.06418 | 0.3385 |
| Site | -0.0009114 | 0.00214 | 0.6703 |
| Motor Control | 0.01964 | 0.02497 | 0.4317 |

**With FSIQ:** F statistic = 5.84, df = 785, adjusted R^2^ = 0.07

| variable | b value | SE | p value |
| --- | --- | --- | --- |
| (Intercept) | -0.4494 | 0.3085 | 0.1455 |
| ADHD-I | -0.02518 | 0.07312 | 0.7306 |
| ADHD-C | -0.05847 | 0.07898 | 0.4594 |
| ASD | 0.1983 | 0.08912 | 0.02635* |
| SLD-Reading | 0.163 | 0.07974 | 0.04124* |
| SLD-Math | 0.02869 | 0.07994 | 0.7198 |
| Anxiety | -0.0005098 | 0.06736 | 0.994 |
| Depression | 0.08449 | 0.1144 | 0.4603 |
| Sex | -0.07237 | 0.06505 | 0.2663 |
| Age | 0.08018 | 0.01346 | 3.86E-09*** |
| SES | -0.03767 | 0.06517 | 0.5634 |
| Site | 0.0001767 | 0.002204 | 0.9361 |
| Motor Control | 0.03876 | 0.0267 | 0.147 |
| WISC-FSIQ | -0.005142 | 0.002579 | 0.04654* |

### **Table S11:** PC2 regressed on dimensional measures

**Without FSIQ:** F statistic = 6.22, df = 783, adjusted R^2^ = 0.06

| variable | b value | SE | p value |
| --- | --- | --- | --- |
| (Intercept) | -1.029 | 0.198 | 2.58E-07*** |
| SWAN-IN | 0.03045 | 0.0329 | 0.3549 |
| SWAN-HY | 0.02651 | 0.03515 | 0.4509 |
| SRS | 0.003108 | 0.005525 | 0.5739 |
| SCQ | -0.00179 | 0.00194 | 0.3566 |
| ASSQ | 0.01676 | 0.008523 | 0.04957* |
| Sex | -0.03367 | 0.0651 | 0.6052 |
| Age | 0.08787 | 0.01327 | 6.60E-11*** |
| SES | -0.0008499 | 0.002177 | 0.6964 |
| Site | -0.05128 | 0.06395 | 0.4229 |
| Motor Control | 0.01984 | 0.02493 | 0.4265 |

**With FSIQ:** F statistic = 6.51, df = 782, adjusted R^2^ = 0.07

| variable | b value | SE | p value |
| --- | --- | --- | --- |
| (Intercept) | -0.4303 | 0.282 | 0.1275 |
| SWAN-IN | 0.01258 | 0.03328 | 0.7057 |
| SWAN-HY | 0.02822 | 0.03498 | 0.4201 |
| SRS | 0.00397 | 0.005505 | 0.471 |
| SCQ | -0.001686 | 0.001931 | 0.3828 |
| ASSQ | 0.01456 | 0.008513 | 0.08756 |
| Sex | -0.0342 | 0.06478 | 0.5977 |
| Age | 0.08793 | 0.0132 | 5.20E-11*** |
| SES | 0.0009451 | 0.002249 | 0.6745 |
| Site | -0.02454 | 0.06427 | 0.7027 |
| Motor Control | 0.04717 | 0.02646 | 0.07505 |
| WISC-FSIQ | -0.00683 | 0.002301 | 0.003087** |
